# Local contribution to the somatosensory evoked potentials in rat’s thalamus

**DOI:** 10.1101/2023.05.25.541803

**Authors:** Władysław Średniawa, Zuzanna Borzymowska, Kacper Kondrakiewicz, Paweł Jurgielewicz, Bartosz Mindur, Paweł Hottowy, Daniel Krzysztof Wójcik, Ewa Kublik

## Abstract

Local Field Potential (LFP), despite its name, often reflects remote activity. Depending on the orientation and synchrony of their sources, both oscillations and more complex waves may passively spread in brain tissue over long distances and be falsely interpreted as local activity at such distant recording site. Current Source Density method was proposed to recover locally active currents from multi-site LFP recordings. Here we use a model-based kernel CSD (kCSD) to study the contribution of local and distant currents to LFP recorded with dense multichannel probes from rat thalamic nuclei and barrel cortex, activated by whisker stimulation. We show that the evoked potential wave seen in the thalamic nuclei around 7–15 ms post-stimulus has a substantial negative component reaching from cortex. This component can be analytically removed and truly local thalamic LFP, with purely thalamic contributions, can be recovered reliably using kCSD. In particular, concurrent recordings from the cortex are not essential for reliable thalamic CSD estimation. Proposed framework can be used to analyse LFP from other brain areas and has consequences for general LFP interpretation and analysis.

**BRIEF SUMMARY:** While recording LFP simultaneously in multiple structures, we often see significant correlations between the observed waves. A natural question is if they are propagated passively from one structure to another or if they are simultaneously generated by different, separated sets of sources. We argue this can be answered reliably using CSD analysis. We focus on the case of thalamic and cortical recordings in the somatosensory system in response to whisker stimulation where we observe significant correlations between early thalamic and cortical responses to whisker deflection.

## 2 INTRODUCTION

The local field potential (LFP) recordings are well suited for both chronic and acute monitoring of neuronal activity at the population level. LFP mainly reflects synaptic and corresponding return currents, although it may also contain contributions from spiking activity, glia, or other sources (Nunez and Srinivasan, 2006; Buzsáki et al., 2012; Einevoll et al., 2013; Głąbska et al., 2017). Due to the volume conduction, high amplitude oscillations in the electric field can propagate widely within the brain tissue. Thus, the signal recorded in one place may have a substantial contribution from distant neuronal populations (Mitzdorf, 1985; Einevoll et al., 2013; Łęski et al., 2010). There is no consensus among researchers on the magnitude of this passive spread. Some reports claim that the recorded potential is almost exclusively a sum of signals originating in a radius about 250 *µ*m from an electrode (Xing et al., 2009; Katzner et al., 2009), while others consider larger distances: Berens et al. (2008) estimated gamma-band spatial resolution to be about 600–1000 *µ*m ; Hunt et al. (2011) found that high frequency (around 150 Hz) oscillations can be detected by electrodes placed several millimeters from their source; we found considerable similarity between LFP collected over the range of about 3 mm in the rat forebrain (Łęski et al., 2007, 2010). Kajikawa and Schroeder (2011) calculated that both lateral and vertical spread of potential at 6 mm from its generator has still 50% of their original magnitude. Kreiman et al. (2006) found that stimulus preferences in visual cortex are similar on a scale of around 5 mm when estimated by the amplitude of evoked field potentials, while for spikes the range of substantial similarity was only around 800 *µ*m. They propose that this difference may be at least partially due to volume conduction. Clearly, the level of LFP correlation between different sites depends on the studied structures, animal model, and applied recording protocol (Lindén et al., 2011).

The theoretical considerations of how far can field potential spread must be confronted with practical considerations addressing the origin of signals recorded from structures of interest. This question is particularity relevant when we record from subcortical nuclei composed of stellate neurons forming nearly closed electric fields. We do record field potential in such structures and they contain a range of physiological oscillations and small but clear waves evoked by sensory stimuli (i.e. evoked potentials). Identification of local and remote origins of these signals and separation of components coming from different anatomically well-defined regions remains an open question.

An important subcortical structure is the thalamus: centrally located mass of neurons clustered in multiple nuclei, with vast network of connection and complex functions. Classical view on thalamus highlighted its role in relaying information to and from neocortex. Currently it is also believed to play a critical role in gating and modulating information transfer not only between periphery and cortex but also between hierarchically organized cortical areas (Sherman and Guillery, 2011). Thalamus and thalamo-cortical loops are believed to be involved in mechanisms responsible for high level cognitive functions (see a review by Wolff and Vann (2019)).

Laboratory rodents offer a convenient model to study the thalamus and thalamo-cortical interaction and LFP is well suited for stable, chronic in vivo monitoring of population level activity. It is, however, essential, though not always easy, to properly understand the components of recorded signals. In our previous experiments we showed in a rat that the early negative (around 10 ms poststimulus, N1_2_) thalamic wave coincides with the strong negative cortical wave (Fig. 1; see also: Kublik et al. (2003); Łęski et al. (2010); Sobolewski et al. (2010)). The amplitude of the thalamic N1_2_ was modulated during cortical cooling — it precisely followed the changes of the amplitude of the cortical potential. We interpreted this as a result of the active influence of the cortical feedback connection over thalamic activity (Kublik et al., 2003; Diamond et al., 1992b). However, in light of our later results (Łęski et al., 2007, 2010) and those from other groups (Kreiman et al., 2006; Kajikawa and Schroeder, 2011), we see that such similarity of thalamic and cortical recordings, to some extent, must be a result of passive spread of the strong cortical signal to the thalamus. Indeed, the cortical representation of mystacial vibrissae in rodents is located relatively close above the somatosensory thalamic nuclei (ventral posteromedial thalamic nuclei, VPM, and medial part of the posterior thalamic nuclear group, PoM). In the rat, the distance between the center of the somatosensory thalamus and the barrel field is around 4 mm and the interval between their borders is around 2 mm (own measurements). In mice, it is about 2.7 mm and 1.4 mm, respectively (estimation based on data from the Allen Mouse Brain Atlas, http://atlas.brain-map.org/). This falls within the distance of the above-mentioned estimates of a field potential passive spread.

**Figure 1.**
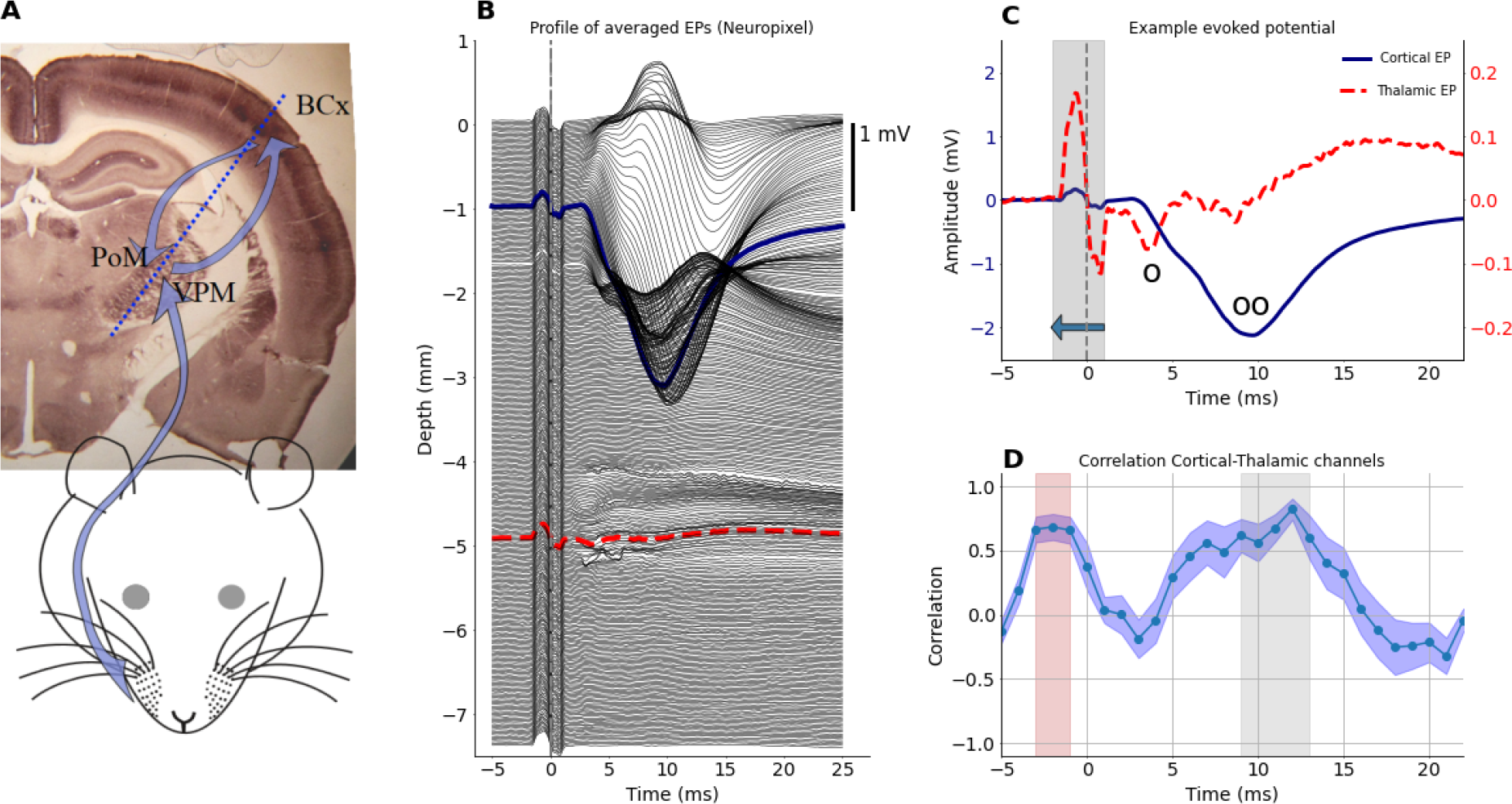
A: Schematic picture of the sensory pathway from the whisker pad to the thalamus (PoM and VPM) and somatosensory barrel cortex (BCx, both clearly visible on cytochrome oxidase stained coronal slice of a rat brain). Blue arrows indicate information flow from periphery to the cortex and a recurrent connection back to the thalamus. In the thalamus, whisker responsive area can be found in the upper-lateral sector of both VPM and PoM nuclei (near arrow heads). Trigeminal ganglion and brain stem nuclei are not marked for clarity of the schema. B: Depth profile of average LFP responses (EPs) to whisker stimulation obtained from Neuropixels inserted at ∼30 deg through the cortex to deep thalamic locations (approximate). Out of 384 Neuropixels channels, uppermost contacts were above the cortical surface while the deepest recording level was around 7.5 mm in the brain. C: Comparison of EPs from a medial cortical channel (indicated by the solid blue line in B) and a thalamic channel (indicated by the dashed red line in B). Vertical dashed line marks the stimulation time (note a bipolar wave of the stimulus artifact around zero, which increases the correlation between thalamic and cortical waveforms presented in D). Circles mark the components of early thalamic negative wave N1_1_ (o) and N1_2_ (oo); the second coincides with strong cortical potential. Note the difference in magnitudes of the cortical and thalamic EPs — left ordinate for cortex, right ordinate for thalamus. D: Average rolling correlation across all experiments (n=11) computed between the central cortical channel and the central thalamic channel (chosen from the histology). Values computed in 3 ms window (represented by gray area in C) were attached at a time point corresponding to the beginning of that window (indicated by the horizontal arrow in C) The blue band along the plot indicates the standard error of the mean across all experiments. Gray shading marks the time when the mean correlation is above 0.5. A similar effect before whisker deflection marked in pink is a consequence of the stimulus artifact.

The main aim of the present study was to determine what part of the field potential recorded in the somatosensory thalamic nuclei is generated locally and what part is a volume-conducted contribution from the primary somatosensory cortex. To examine this problem, we recorded evoked potentials (EPs) from multiple locations through the barrel cortex and thalamus, and we applied kernel Current Source Density method (kCSD, Potworowski et al. (2012); Chintaluri et al. (2021)) to estimate the distribution of current sinks and sources along the shaft of the electrode (Mitzdorf, 1985). Next, we used them, in a forward modeling approach (Einevoll et al., 2013), to compute contributions coming from thalamic or cortical areas to the potential measured at different electrode locations.

## 3 RESULTS

### 3.1 LFP in the barrel cortex and thalamic area

We recorded whisker-evoked field potentials in the primary somatosensory cortex and in the somatosensory thalamus of the rat’s right hemisphere (Fig. 1A) and repeated this paradigm with three different experimental setups (see Methods). Example results for simultaneous thalamo-cortical recordings done with a 384 channel Neuropixels probe are presented in Fig. 1B. Note that during the strongest negative wave recorded in the cortex, around 10 ms poststimulus (see ‘oo’ mark in Fig. 1C), a negative deflection of the potential recorded in the thalamus occurs. To quantify similarity of thalamic and cortical EPs, in each rat we computed their correlation in a 3 ms window rolling along EP waveforms (Fig. 1D). For a group of 11 rats, the correlation value above 0.5 was observed around 10–12 ms after the stimulus.

Similarity of the cortical and thalamic negative waves around 10 ms suggests that they may reflect partly the same currents, which — observed from different distances — result in waves of larger (cortex) or smaller (thalamus) amplitudes. However, the amplitude and polarity of evoked potentials change along the depth profile spanning from cortex to the thalamus. The largest wave around 10 ms post stimulus is the negative deflection in the cortical middle layers; but at the same latency, the potentials at the bottom of the cortex have a clear positive polarity (Fig. 1B, 2A). Interestingly, the polarity of the thalamic recordings remains negative across a range of depths which seems consistent with the cortical signal recorded in the middle layers rather than with the positive signal recorded deeper.

To better understand the relation between these cortical and thalamic signals we made a simple estimation of the potential spread: we calculated currents from EP profile and selected cortical CSD channels (example in Fig. 2B). We then estimated field potentials that these cortical sources would induce in the tissue along the electrode line going from brain surface into the thalamus. As can be seen in Fig. 2C, the strong negative cortical field has a very long range. The purple stripe around 10 ms is only transiently reversed at the bottom of the cortex, and it is evident in a whole sub-cortical space below. This example highlights the strength of passive spread (i.e. volume conduction) from cortical currents and their possible impact on distal (thalamic) potential values.

**Figure 2.**
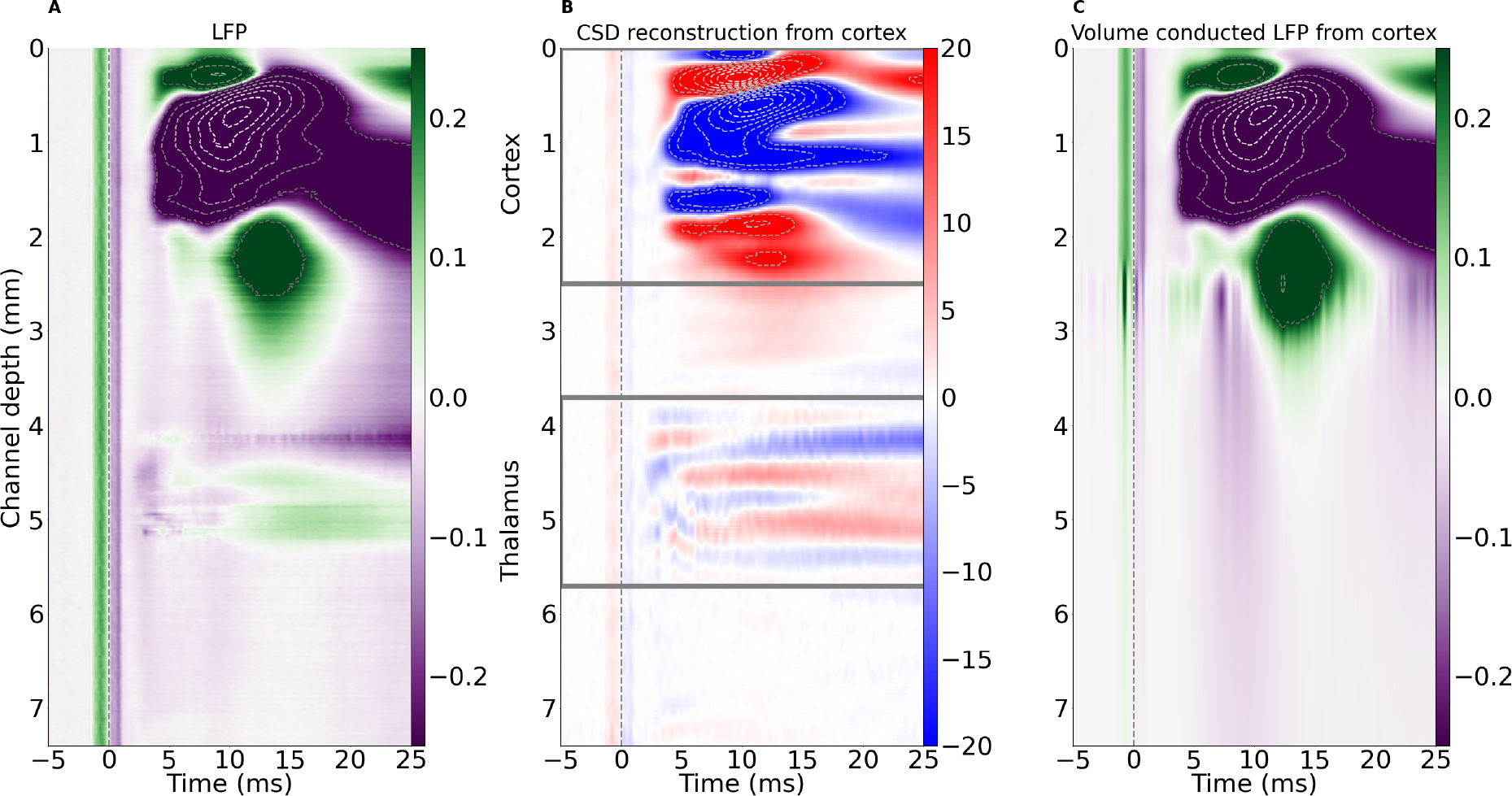
A: Example spatiotemporal EP profile from Neuropixels probe (same data as in Fig. 1B) presented as a 3D color map. B: Spatiotemporal CSD profile estimated from data in A. Note evident current sinks and sources in the thalamic region. C: Spatiotemporal profile of EPs estimated in the whole cortico-thalamic space from recordings restricted to cortical area marked in B by a gray rectangle. Note a stripe of negative (purple) potential around 10 ms spreading from cortical sources to deep subcortical levels, which may modify thalamic waveforms. In all panels, horizontal axis shows the time with respect to the stimulus (applied at t=0); left vertical axis shows recording depth; colorbars on the right of each panel show the magnitude of potential (A and C, [mV]) and current source density (B, [*µ*A/mm^3^]). Vertical dotted lines at time 0 mark the stimulus onset and related artefact in the data.

We thus stated a hypothesis that a strong activity from middle cortical layers can be seen even 3 mm from the source and can overshadow subcortical signals, and we propose a general CSD-based pipeline to recover the local contributions to field activity. This analytical pipeline is of general validity even if here it is illustrated with examples from the primary somatosensory pathway.

### 3.2 CSD reconstruction and forward modeling

Generally used approach to extract local activity from LFP is the CSD method. We used kCSD method, which enables reconstruction from nonlinearly distributed electrode contacts and in arbitrary space of sources. The example CSD computed from the recordings from Neuropixels probe (Fig. 2A) is plotted as a spatio-temporal profile along electrode track (Fig. 2B). As seen in case of potential values the density of currents in the cortex is much higher than in thalamus. However, the analysis revealed clear local thalamic activity not only at the earliest post-stimulus time (before evolution of cortical activity), but also later, dipole-like current sinks and sources.

Having estimated the CSD, we can compute how much a given subset of sources contributes to potentials at each recording site. This should show us how thalamic LFPs would look with and without a cortical component and vice versa (Fig. 3).

**Figure 3.**
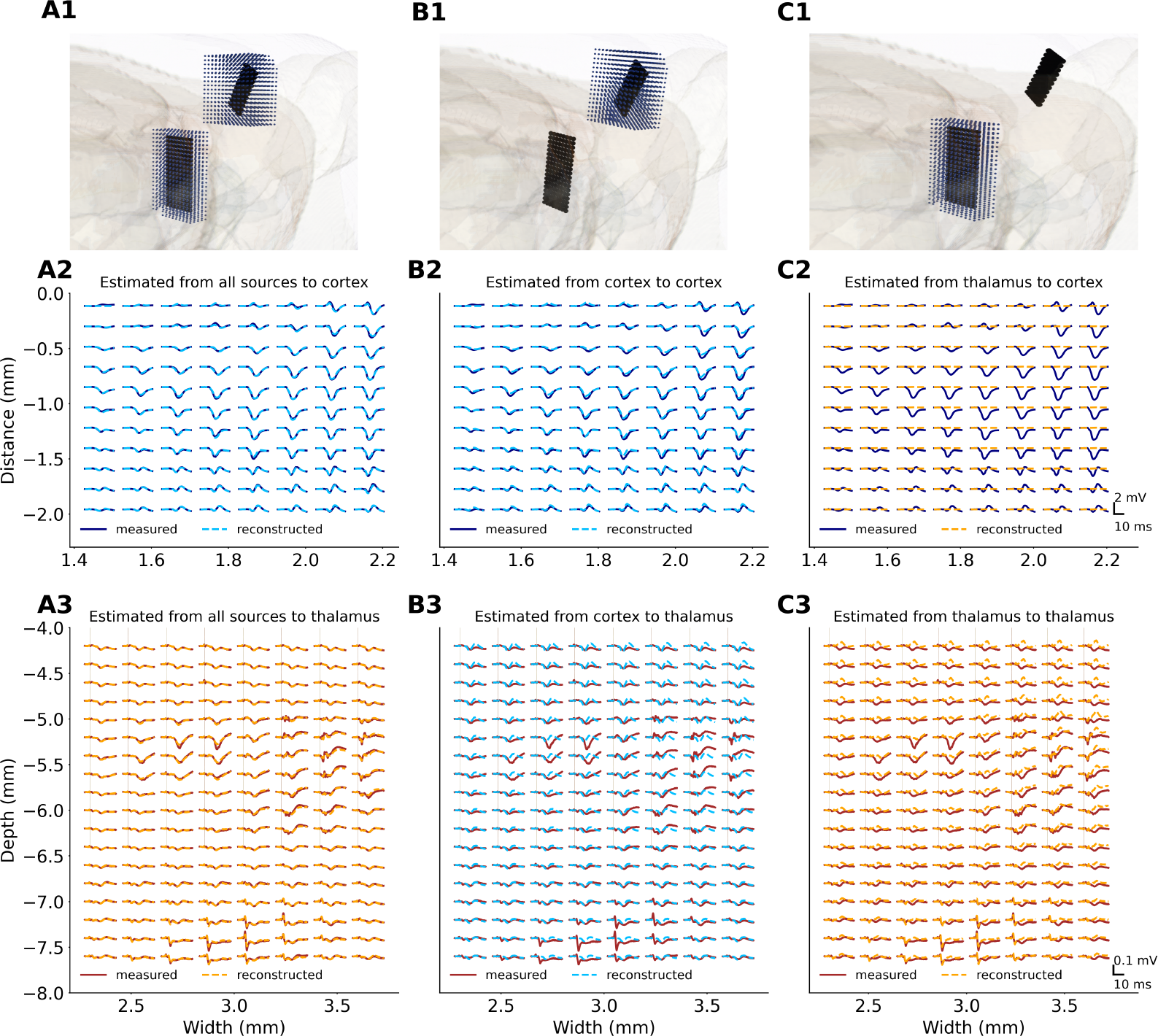
Comparison of measured EPs with their counterparts computed from subsets of current sources (example from an experiment using 8x8 Neuronexus probes). Panels in the first row (A1, B1 and C1) show schematic representations of sources reconstruction space (black dots: electrode positions; blue dots: subset of CSD used for computation of cortical (B) and thalamic (C) contributions to the EPs. Measured EPs and computed contributions are overlaid in the second (for cortical channels) and third row (for thalamic channels). Note a lateral gradient of EP amplitude in the cortex — NeuroNexus A8x8 silicon probe was inserted on the edge of barrel field (see Suppl. Fig. 2B). In the thalamus (A3, B3, C3) we can spot two clusters of early evoked responses — one (middle-right area) around the main whisker representation in VPM/PoM complex, the other (lower edge) corresponding to zona incerta nucleus. (A) All the cortical and thalamic sources were used for computed EPs shown in A2, A3 panels. Obtained EPs are consistent with the measurements which shows the method is self-consistent. (B) Cortical contributions to the measured EPs (B2, B3). Estimated potentials fit well cortical measurements (B2) and show similarities with thalamic recordings (B3). (C) Thalamic contributions to the measured EPs (C2, C3). As expected, the thalamic contributions to the cortical channels are negligible. On the other hand, the truly local part of the thalamic LFP, which is that estimated from thalamic sources (dashed orange line), is different from the measured EPs, which are contaminated by passively propagated strong cortical signal (dashed blue line in B3).

### 3.3 LFP reconstruction from subset of sources

In all datasets we used CSD reconstructed from measured EPs to estimate potential waveforms, first from full CSD profiles, and then separately from its cortical and thalamic subsets. The results are illustrated in Fig. 3 with an example from one of 2D EP profiles obtained with A8x8 NeuroNexus probe. Taking all the sources for reconstruction we obtained LFPs almost identical with measured ones (Fig. 3A1–A3; as required by self-consistent properties of kCSD). Cortical recordings and their reconstructions from cortical-only sources mostly overlapped (Fig. 3B2) which indicates that cortical potential has indeed mainly local origin. The influence of thalamic sources was minimal in the cortex — the reconstructed contributions from thalamic sources to cortical potential were practically flat (Fig. 3C2).

This is not the case in the thalamus. Here, the locally generated part (estimated from thalamic sources, Fig. 3C3, dashed orange) is much closer to the measured potentials than the cortex based estimation (Fig. 3B3, dashed blue) but on average more positive. Cortical contributions to thalamic evoked potentials constituted relatively large negative wave all over the thalamic space (Fig. 3B3). Clearly, the LFP recorded in the thalamus contains a mix of thalamic and cortical contributions each of which must be recovered with the help of CSD methods.

### 3.4 Estimating overall effect of cortical LFP volume conduction

Fig. 4A shows enlarged overlay of exemplary measured and estimated, cortical and thalamic, EPs (examples from Fig. 3B2 and C3). Contributions from cortical sources were sufficient to fully reconstruct cortical EPs, while thalamic sources were not sufficient to precisely reconstruct thalamic EP. Contributions from cortical sources reliably reconstructed cortical EPs, while reconstruction from thalamic sources differed from the measured thalamic EPs. Real EP trace tended to be more negative, in particular from around 6 to 14 ms, than the reconstructed ones (see Fig. 4B and C for a group summary of EP amplitute at 10 ms post stimulus, B: permutation paired-test, p-value=0.002, n=11, C thalamic channels: 1 sample ttest, p-value=0.013, C cortical channels: 1 sample ttest, p-value=0.44). We argue this is a consequence of the strong negative wave conducted from the cortex.

**Figure 4.**
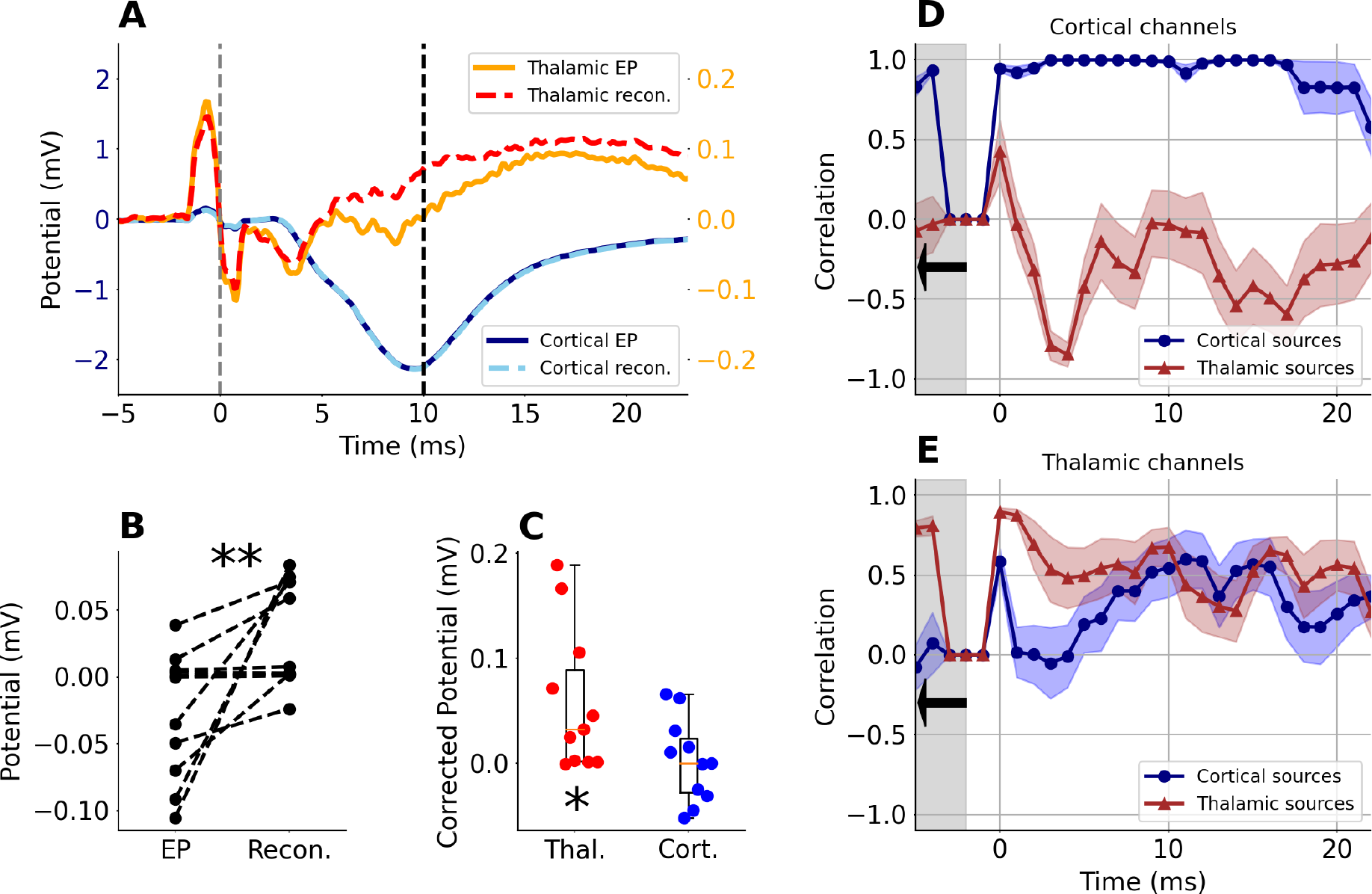
A: Examples of measured and reconstructed potential from central cortical (cyan and dotted-blue, left Y axis) and thalamic sites (orange and red-dashed, right Y axis) from Figure 3 B2 and C3 panels. Solid lines represent measured potentials, dashed lines represent reconstructed potential from local sources. B: Potential values at 10 ms after whisker stimulation in recorded thalamic EPs and their reconstructions. Note significantly less negative values in reconstructed data (permutation paired-test, p-value=0.002, n=11. C: Absolute reconstructed-minus-measured values for thalamic (red) and cortical (blue) potential wave at 10 ms post-stimulus. Note that the difference is significantly different from zero in thalamus but not in the cortex. (reconstructed-measured) in thalamic area (orange) and in cortical area (blue). There is a significant difference from 0 in the thalamic area, (1 sample ttest, t=3.03, p-value=0.013, n=11) but not in the cortex (1 sample ttest, t=0.8, p-value=0.44, n=11) D: Average correlation score in cortical channels between measured EPs and EPs reconstructed from cortical (blue) or thalamic (red) sources. E: The same as B but for thalamic channels. Shaded corridor along line-plots in B and C represent SEM (n=11 rats).

To quantify the similarities and differences between the waveforms of measured EPs and reconstructed contributions from thalamic and cortical sources we calculated their correlations in a 3 ms window running along the EP traces. In each rat, correlation coefficients were computed separately for the thalamus and for the cortex for the channels that were identified in histological data as located in the centers of analysed structures (details in Suppl. Table 1). Group averaged results (n=11 rats) are presented in Fig. 4D and E. The correlation score for the cortical channels (Fig. 4D) was close to one throughout the analyzed time (25 ms) when reconstructed from cortical sources and it was close to zero or negative when comparing measurements with contributions to cortical potentials from thalamic sources.

Correlation coefficient for thalamic waveforms measured and reconstructed from thalamic sources (Fig. 4E, red line) was the highest (0.87±0.05) in the early post stimulus window, fell down after 5 ms and was close to zero (0.21±0.14) at 14 ms (Fig. 4E, red line). Similarity of the real thalamic EPs to those estimated from cortical sources showed an opposite pattern (Fig. 4E, blue line). Correlation was close to zero in an early window up to 5 ms post stimulus but then increased reaching a maximum up to 0.59±0.18 around 11 ms and then dropped again.

This group analysis confirmed that strong cortical currents indeed may strongly affect the field potentials recorded in the thalamus. This influence was maximal in a time window between 10 and 15 ms, when LFP recorded from thalamus was defined predominantly by cortical sources.

### 3.5 Detecting weak thalamic activity in the CSD space

Our analyses showed that thalamic LFP can be substantially contaminated by passive contributions from strong sources, cortical in this case, and that CSD reconstruction allows to identify truly local contributions. This observation leads to a range of practical questions: how to reconstruct this local activity most reliably? Where to locate the electrodes? Should one record from all the possible (close and far) current sources, or is it enough to monitor the local activity? How to optimally choose CSD reconstruction space in a model-based reconstruction method such as kCSD?

To address these issues we considered four analytical setups using data from an experiment with two independent A8x8 silicon probes (Fig. 5). In consecutive analyses we assumed that either (1) only the thalamic electrode was used, probing relatively weak thalamic responses, or (2) both electrodes were used, one in thalamus and the other in the cortex, probing the stronger cortical activity. The reconstruction was performed in two different source spaces: one, limited to thalamic region and the other, covering substantial part of the forebrain including all thalamic and cortical electrodes. We then estimated CSD in four available combinations of electrode setups and CSD spaces (Fig. 5, top row).

**Figure 5.**
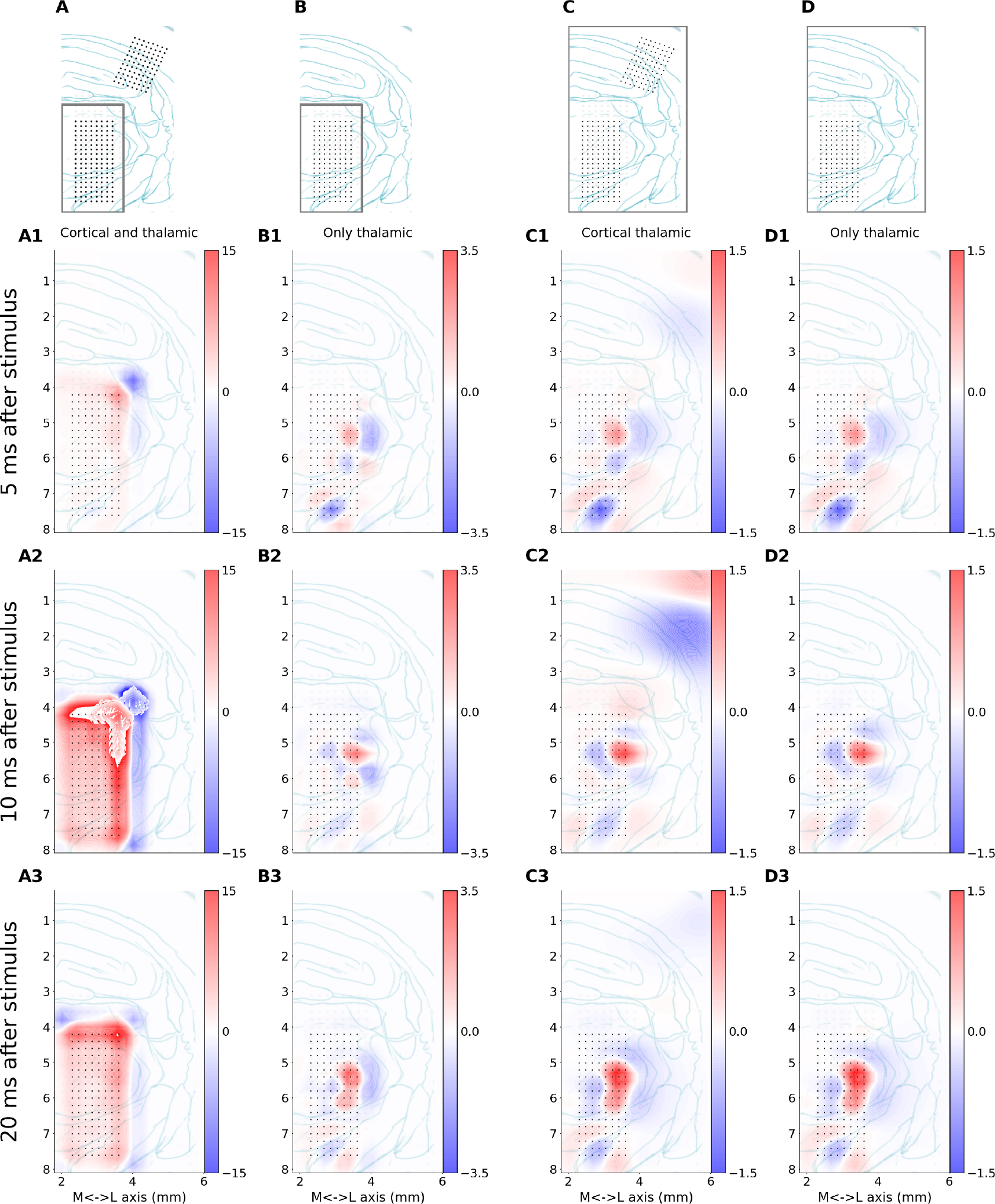
The first row shows the analytical setup used in the analysis in a given column. The dots represent the electrodes taken into consideration. The frame indicates area where sources were assumed (basis sources were placed). A: Recordings from the thalamus and cortex considered; sources assumed only in the thalamus. B: Only thalamic recordings considered; sources assumed only in the thalamus. C: Recordings from the thalamus and cortex considered; reconstruction in a large block of tissue covering thalamus and cortex (same as in D). D: Only thalamic recordings considered; reconstruction in a large block of tissue covering thalamus and cortex (same as in C). Rows A1–A3 show CSD reconstructions snapshots at 5, 10 and 20 ms after stimulus that were analyzed assuming setups indicated in the first row. CSD spatial maps are overlaid on a drawing representing histological verification of recording points (dots) location within thalamic structures. X and Y axes are scaled in millimeters with horizontal zero at cortical surface above thalamus and lateral (X axis) values measured from mid-line. Coronal plane corresponds to ∼ 3 mm posterior from bregma point. Note the different color scales shown next to each panel.

Of these four analysis three rendered consistent results (Fig. 5B–D), the other was erroneous (Fig. 5A)c— where we tried to explain simultaneously strong cortical and weak thalamic recordings with sources placed only in the thalamus. As long as the reconstruction space encompassed all the electrode positions, obtained CSD reconstructions seemed credible. What we consider natural approaches to CSD analysis of these data are shown in column Fig. 5B, where the reconstruction space is a region in the thalamus containing the electrodes with a 0.5 mm margin to accommodate possible artifacts from remote sources, and in Fig. 5C, where the information is taken from both thalamic and cortical electrodes and reconstruction space covers all electrode positions. Using only thalamic data for reconstruction in the larger space does not significantly affect thalamic reconstruction Fig. 5D. However, keeping reconstruction space restricted to the thalamus and demanding it to explain both thalamic and cortical recordings requires the method to compensate the missing cortical sources present in Fig. 5C with a strong non-physiological dipolar halo surrounding the thalamic electrodes (Fig. 5A), which could not be aligned with the anatomy of thalamic nuclei, and which is a clear artifact.

The results shown in Fig. 5B–D provide similar patterns of detailed thalamic currents, clearly and reasonably aligned with the histology. Clusters of earliest sinks and sources (5 ms, Fig. 5A1–D1) were localized precisely in the regions receiving direct peripheral input — the dorsal sector of VPM and PoM, and zona incerta. Importantly, this well localized pattern was preserved also in later post-stimulus windows (see Fig. 5, rows 2–3), which in the LFP picture would be overflown by the strong cortical negative wave. The main effect of the extension of estimation space from Fig. 5B to D was a reduction in magnitude of thalamic currents (compare colour intensity between B and C–D in Fig. 5), however the source pattern was the same. Interestingly, the inclusion of cortical signals in estimation (Fig. 5D) did not significantly affect the pattern of reconstructed CSD in the thalamus, i.e. it did not improve their separation. This is what we expect from traditional approaches to CSD estimation using three-point second derivative Pitts (1952) and it is also true here for this model-based analysis.

## 4 DISCUSSION

### 4.1 Summary

In the current paper, we have addressed the problem of ambiguous origin of subcortical local field potentials. We recorded simultaneous whisker evoked responses from somatosensory thalamus and cortex in rat. We confirmed that the earliest waves in thalamic LFP are of local origin (they correspond to the activation by peripheral input) but strong impact of cortical field is passively shaping thalamic signal in consecutive (7–15 ms) poststimulus time window. Therefore, it is necessary to precede the interpretation of such data with appropriate analyses. We showed that with use of kernel CSD analysis it is possible to recover the contribution of weak local currents to the thalamic waves.

Extracellular electrophysiological recordings reflect activity of local and distant generators. Mixing of such varied contributions makes it difficult to interpret the data, especially in the case of closely located and interconnected structures, where waves resulting from real interactions are entangled with volume conducted ones. This reservation may apply to any field recordings in animals as small as laboratory rodents, however, it has also been raised for signals from human intracranial electrodes (Wennberg and Lozano, 2003). Our example comes from a rat vibrisae-barrel system which is an important experimental model. Thanks to its easily accessible receptive field (whisker pad) and large and clear somatotopic representation (Welker, 1976; Diamond et al., 1992a; Land et al., 1995; Fox and Woolsey, 2008) it is widely used to study neuronal plasticity (Erzurumlu and Gaspar, 2020) but also information processing within feed-forward and reciprocal thalamo-cortical connections (Petersen, 2007; Sobolewski et al., 2010, 2015). In anatomical studies, the large size of cortical barrel fields is an advantage. However, it is not so beneficial in case of in vivo recording of electrical activity outside barrel cortex. In the thalamus the earliest responses to whisker stimulation start around 3 ms post stimulus. At this time single or multi-unit activity can be observed followed by a small-amplitute negative potential wave peaking around 8–12 ms (e.g. Fig. 1C). Whisker evoked response in the cortex starts from ∼5 ms post-stimulus (Armstrong-James, 1995) and it develops a very large negative wave which peaks, at the level of layer 4/5, with a latency of around 10 ms. Here we showed that the trace of this field, generated in the large volume of active barrel cortex, can affect recordings in subcortical structures including thalamus. There are also examples that barrel cortex activity may reach less obvious locations like the olfactory bulb (Parabucki and Lampl, 2017). Another substantial structure whose activity may spread widely in a brain is hippocampal formation. It is believed to oscillate in synchrony (in the theta rhythm) with other nodes of its functional network (Nuñez and Buño, 2021). However, it has recently been estimated, at the level of field potential recordings, that some of these synchronizations are illusory: theta oscillation in the habenular LFP signals (Goutagny et al., 2013) came out not to be locally generated in synchrony with hippocampus but simply passively conducted from it (Bertone-Cueto et al., 2020). Similarly, synchrony of high-frequency oscillations (HFO) between hippocampus and nucleus accumbens was shown to be a consequence of passive field propagation (Hunt et al., 2011).

Cortex and hippocampus are laminar structures with multiple parallel pyramidal cells forming synchronous electrical dipoles upon synaptic activation and generating open electric fields, which can be recorded even from far away. Many subcortical nuclei are composed of stellate neurons with roughly spherically symmetric dendritic trees (never perfectly symmetrical) and closed electric fields which are much more difficult to detect, in particular from a distance. Modelling indicated that the electrodes penetrating such a structure should reliably record its local signals (Tanaka and Nakamura, 2019). Experimental data confirmed that subcortical LFP is built by well-defined local sinks and sources (Łęski et al. (2007, 2010) and current data), however, it is contaminated with distant influences.

### 4.2 Extracellular electrophysiology versus other techniques

Classical electrophysiology is not the only technique to monitor neuronal electrical activity and ionic flow. Recent development of optical imaging (i.e. new, faster voltage sensitive and calcium indicators, and multi-photon microscopy (Antic et al., 2016) enabled monitoring electrical events in tissue with good spatial resolution and without the burden of volume conduction, electromagnetic interference and ambiguous referencing. The methodology is widely applied for little-invasive studies of the brain cortex, and the use of implantable optical fibres or gradient-index (GRIN) lenses pushes the imaging plane to subcortical level (see review by Zhang et al. (2019)). However, due to their large diameter (∼1 mm), GRIN lenses cause tremendous trauma to the brain while multichannel silicon probes, with their small diameter (∼ 100 *µm*×20 *µm*), are much less invasive and preserve neural tissue in a more physiological state. Also, the time resolution of optical imaging techniques is lower than in electrophysiology. Voltage sensitive indicators used to record whisker evoked activity in mouse somatosensory thalamus (Tang et al., 2015) completely overlooked earliest responses. Peripheral input evokes spiking activity with a few millisecond latency (Armstrong-James and Callahan, 1991; Diamond et al., 1992b), while the optical thalamic response reported by Tang et al. (2015) started after 20 ms post-stimulus. Stronger, cortical optical responses were noticeable, as expected from electrophysiology (Armstrong-James and Callahan (1991)), in an early window between 5 and 10 ms post-stimulus. Thus, electrophysiological recordings using implanted multichannel probes remain the simplest and least invasive tool for detecting neuronal activity, including relatively weak signals, from multiple subcortical sites in the brain. Silicon probes allow recording from multiple structures along one shaft, can be reinserted multiple times during one surgery, and 3D patterns of multiple shafts can be chosen or designed to fit anatomical requirements of particular experiments. As we have shown in this report, appropriate analyses help overcome the difficulties posed by volume conductivity.

### 4.3 Field potentials versus spikes: complementary, not an alternative

Wideband extracellular signals can be broadly divided into low and high frequency components. The Low Frequency Part, as we can also spell out the LFP (Głąbska et al., 2017), is built up mostly by slow transmembrane currents, while the latter includes traces of action potentials (spikes) fired by single or multiple neurons in the vicinity of a microelectrode tip (Destexhe and Bedard, 2013). Spikes’ amplitudes drop quickly with increasing distance from an electrode and the size of an electrode (Moffitt and McIntyre, 2005; Pettersen and Einevoll, 2008). Thus, in case of spikes, there is little or no confusion regarding the location of their generators. While the location of LFP remains a challenge, there are multiple advantages of field potential recordings and analysis (Buzsáki et al. (2012); Herreras (2016); Jackson and Hall (2017)). Spikes and LFP carry different information about neuronal activity and cannot be used interchangeably. Slow postsynaptic potentials dominating LFP inform about the input to the neurons, the analogue processes of data accumulation and analysis realized on neuronal membranes through summation of postsynaptic depolarizations and hyperpolarizations. This may or may not lead to generation of an action potential depending on whether the spiking threshold is reached. Spiking activity is thus interpreted as an output of the system. These different roles of the two communication channels can be utilized in analysis, for instance Laminar Population Analysis attempts to establish functional connectivity pattern within a network combining the information about the inputs and outputs from multiple locations in a structure (Głąbska et al., 2016). This is why it is best to record wideband signal and analyze both action and field potentials. However, this may not be possible if an amplifier input range and a bit-depth are not good enough to cover frequency and amplitude ranges of both signal types. We then have to set filtering and choose which component to record and which to discard. LFP is chosen if we are interested in population, subtreshold oscillatory activity. There is also a practical aspect to LFP recording. Implanted probes provoke scarring in neuronal tissue which, in particular in long-lasting, chronic experiments, degrades the quality of spike registration. Currently, great research effort is being put into improving the biocompatibility of implanted probes (Fattahi et al., 2014; Szostak et al., 2017), but in the meantime, LFP is a more stable and predictable signal, in particular in the chronic applications (Jackson and Hall, 2017). It is thus crucial to systematically study the sources of LFP (Herreras, 2016).

### 4.4 Identification and extraction of local and distant contributions to the LFP with experimental interventions

Having decided that using LFP is a viable approach to study subcortical activity we can use experimental and analytical tools to distinguish local from remote contributions. Experimentally we can switch off efferent pathways by cooling, dissection, chemo- and optogenetics. These methods can help identify different contributions but all of them affect the network so in consequence we study an altered system. If we wish to understand minimally affected system there is no alternative to analytical extraction of signal sub-components form physiological in vivo data. Following experimental interventions can help to verify the conclusions regarding their anatomical sources. Surgical interventions can be used to physically separate structures and identify local and distant, active and passive signal elements. For example, dissection of the olfactory bulb blocked the widespread, ketamine-induced high frequency oscillations and confirmed it as a site of their neuronal generator (Średniawa et al., 2021). Whisker evoked field potentials were still recorded in the olfactory bulb after transsection of neuronal pathways, which indicated that they were transmitted passively through touching blocs of cut tissue (Parabucki and Lampl, 2017). While the olfactory bulb is relatively easy to isolate from the rest of the brain, in vivo dissection of structures like the cortex and thalamus is not as feasible. Moreover, brain trauma is huge and effects are not reversible. A less invasive approach involves silencing of chosen structures or neuronal population with physical (e.g. cold), genetic (e.g. opto- or chemogenetics) or pharmacological tools (e.g. lidocaine, tetrodotoxin, magnesium solution). The inactivated structure does not generate its own field potentials, so any locally recorded signal can be interpreted as originating from other generators; also, inactivated structure cannot influence distant regions, neither passively nor actively. Silencing is usually used to study the active synaptic interactions, just like in the case of research showing how the whisker evoked spiking in thalamic sensory nuclei is independent (VPM) or dependent (PoM) on the cortical activity (Diamond et al., 1992a; Mease et al., 2016). On the other hand, we showed that, unlike action potentials, LFP responses in both nuclei similarly followed the changes in cortical activity modulated by a gradual cooling (Kublik et al., 2003). Those results suggested that both thalamic nuclei receive subthreshold cortical input (presumably from layer 6) that can be detected by LFP, but not single unit recordings. However, some of these thalamic potential waves might be a result of volume conduction, not synaptic activation from the cortex. This reservation was confirmed by the current analyses. CSD and modeling approach allowed to exclude the cortical generators and reconstruct “pure” thalamic EP in which the negative deflection around 10 ms was substantially reduced as compared to the LFP (Fig. 4 A–C). We further confirmed this analytical approach with experimental intervention: we repeated whisker stimulation and kCSD analysis after topical lidocaine application over barrel cortex (see Supplementary materials). Thalamic EPs, recorded during cortical inactivation, were no longer significantly different from reconstructed EPs (Supp. Fig. 1 D, E) but were slightly different from the control ones (Supp. Fig. 1 D). The difference between recorded thalamic EPs before and after cortical inactivation can be explained by the elimination of synaptic cortico-thalamic interaction. Layer 6 is a well known modulator of thalamic activity (Wróbel et al., 1998; Thomson, 2010; Mease et al., 2014); also layer 5 can induce strong EPSPs in PoM neurons (Mease et al., 2016). In the context of current research, more important was the fact that the lidocaine eliminated cortical generators and volume conducted components from thalamic recordings. Local thalamic currents evoked by peripheral input were the only generators of thalamic LFP and our kCSD analysis was able to reliably reconstruct this effect, which confirms the validity of its results also in control conditions.

### 4.5 Analytical approaches alternative to CSD

Many approaches have been used to analyze relations between continuous signals recorded in different structures (Kamiński and Blinowska, 1991; Wójcik et al., 2001; Kuś et al., 2004; Stam, 2005; Łęski and Wójcik, 2008; Sobolewski et al., 2010) and to untangle the signals comming from different sources (Makeig et al., 1997; Musial et al., 1998; Makarov et al., 2010; Whitmore and Lin, 2016; Bertone-Cueto et al., 2020). We can distinguish approaches grounded in nonlinear dynamics, spectral analysis, signal decomposition, source separation, source reconstruction (CSD analysis) with multiple approaches to functional connectivity. In our view, to obtain actual localization of a given activity there is no alternative to current source reconstruction. With all the limitations of CSD analysis it remains the only approach indicating locality of the signal, effectively deblurring the recorded LFP. While other methods mentioned can help answer questions of signal interactions, indicate common drivers or separate functional components, as long as they are based on raw LFPs the inherent mixing of local and remote sources within each signal is unavoidable and present.

### 4.6 Practical aspects of CSD estimation

The final question remains how to best estimate the distribution of current sources from a given set of recordings obtained in a given anatomical context.

Traditional approach of computing three-point approximation to the second derivative was proposed by Pitts (1952), who calculated a map of spinal cord activity from 2 D grid of recording points. This method gained popularity after rise of one dimentional, laminar probes, when Nicholson and Freeman (1975) advocated its use in laminar structures, see review by Mitzdorf (1985). This traditional CSD has the advantage of simplicity but numerous disadvantages. Notably, it does not give independent results at the boundary, it requires regular electrode grids, and it gives estimation only at the recording points. These limitations can be circumvented, e.g. with interpolation, but in our view the simplest approach to obtain the distribution of CSD is with model-based methods such as inverse CSD (Pettersen et al., 2006; Łęski et al., 2007, 2011), kernel CSD (Potworowski et al., 2012; Chintaluri et al., 2021), or Gaussian process CSD (Klein et al., 2021). These approaches overcome the limitations of the traditional approaches, allow for self-consistent estimation of potential and CSD at arbitrary points, but since they are model-based it may be less apparent how to best use them. One of our goals here was to discuss the main aspects of model-based CSD analysis in the context of the present data. We focus on kernel CSD of which inverse CSD is a special case (Potworowski et al., 2012).

To apply kernel CSD we first cover the region of interest where dominating sources are expected with a rich family of basis sources, typically small overlapping gaussians covering the region densely. This procedure separates the experimental setup (electrode placement) from analytical setup (estimation space). The number of basis sources is not very important. The more the better but we observe stability with growing number of basis sources, also reasonable independence from the basis source specific shape and size. It is always good to check if significant improvement is obtained for a given analysis when increasing the number of basis sources or modifying their size.

In principle it is possible to estimate CSD in one place (e.g. thalamus) from measurements elsewhere (e.g. cortex), see (Chintaluri et al., 2021) for an example and discussion. In practice, common sense is recommended. The level of noise in the system obviously precludes certain analysis; it is best to record in the system of interest as close to the generating sources as possible. Thanks to separation of the experimental and analytical setups in kCSD, the grids of recording points can be arbitrary, irregular. Electrodes can cluster in distant structures (e.g. cortex and thalamus), probes can have semilinear, zigzag pattern (as in Neuropixels) or broken contacts (as practically each probe in our experiment) and kCSD still gives reliable results.

The price we pay for the flexibility of kCSD is that by its nature the estimation can be obtained only in the space defined by the basis sources. If our initial placement makes no sense, the results of the analysis make no sense. In practice, we find that distributing the basis sources to cover densely the region span by the electrodes gives results consistent with the simplest CSD estimation, that is we effectively compute the derivative of the potential, except we obtain results independent of electrode distribution, smooth, and self-consistent. To compensate for the sources outside the region span by the electrodes we recommend extending the estimation region. In this way the artifacts introduced by the method trying to compensate for the sources beyond our electrodes are typically crammed within the added margin area. See Methods section for the description of our procedure in the present case, e.g. for Fig. 5.

Possible sanity checks of the assumed procedure (combination of electrode setup and basis source placement) are to compare the obtained results with histology or to compute the eigensources for the system (Chintaluri et al., 2021). Comparison against histology, such as shown here in Fig. 5, allows to verify if the obtained results make anatomical sense, e.g. sinks and sources observed in meanigful nuclei or cortical regions. The eigensources are profiles of CSD which can in principle be ideally reconstructed. They span the space of all possible reconstructions that can be obtained for a given combination of electrode distribution and base sources placement.

Noise is another factor we must consider in the analysis. To avoid overfitting to noise we regularize estimation (see Methods). We select the regularization parameter *λ* using cross-validation or the L-curve (Chintaluri et al., 2019). However, when the automatic selection procedure provide unrealistically small *λ* values it is prudent to increase it to further dampen the noise.

CSD reconstruction effectively plays the role of deblurring of the potential landscape. A problem it does not solve is separation of activity of overlapping cell populations. In our experience combining CSD analysis with spatial ICA gives excellent, interpretable results (Łęski et al., 2010; Głąbska et al., 2014) facilitating structural analysis.

In the analysis here we assumed constant, isotropic and homogeneous conductivity. While there are reports of tissue conductivity, in particular inhomogeneity and anisotropy in the cortex (Goto et al., 2010), in our view the variability between different specimen as reported there is so big, that assuming literature values rather than those characterizing the specific animal currently studied we would make bigger estimation errors than simply assuming constant values. This is supported by our finite element modeling in a slice (Ness et al., 2015). The situation is different for inverse methods for EEG or ECoG, where significant changes of conductivity between scalp, skull and brain tissue cannot be ignored, but for recordings deep within tissue we recommend assuming constant conductivity.

CSD has most commonly been used to analyze and interpret LFP profiles recorded from laminar structures with clear dipoles building the signal (Nicholson and Freeman, 1975; Mitzdorf, 1985). However, CSD has also been effective in case of non-laminar structures (Łęski et al., 2007, 2010). While having experimental setup which probes in 2d or 3d gives better insight into the spatial organization of the sources (Pitts, 1952; Łęski et al., 2007, 2010, 2011), even laminar probes give meaningful results, consistent with cuts through higher-dimensional reconstructions, and can fruitfully be used.

### 4.7 Take home message

LFP recordings from rat’s somatosensory thalamus are contaminated by cortical electric field, in particular during strong, whisker evoked excitation. With kernel Current Source Density method we can reliably estimate currents and reconstruct thalamic activity even without monitoring cortical field potentials with additional sets of electrodes. The proposed workflow can be used to identify local contributions to LFP recorded in other subcortical structures.

## 5 METHODS AND MATERIALS

### 5.1 Animals, surgery and histology

The experiments were performed on 11 adult male Wistar rats weighing 350–560 g. All experimental procedures followed EU directives 86/609/EEC and 2010/63/EU and were accepted by the 1st Warsaw Local Ethics Committee. Animals were anaesthetized with urethane (1.5 mg/kg, i.p., with 10% of the original dose added when necessary) and placed in stereotaxic apparatus (Narishige). Local anesthetic (Emla, 2.5% cream, AstraZeneca) was applied to the rats’ ears and the skin over the skull was injected with lidocaine (Lignocainum hydrochloricum 1%, Polfa Warszawa S.A.) prior to the surgery. The skull was opened above the barrel field and the thalamus in the right hemisphere. Fluid requirements were fulfilled by s.c. injections of 0.9% NaCl, 5% glucose, or both. The body temperature was kept at 37–38°C by a thermostatic blanket. To control the physiological condition of the animal heart rate, breath rate and oxygen saturation were monitored during the whole experiment (using MouseOx, Starr Life Sciences Corp.).

After completion of an experiment, rats received an overdose of pentobarbital (150 mg/kg i.p.) and were perfused transcardially with phosphate buffered saline (PBS) followed by 10% formalin in PBS. The brains were removed and cryoprotected in 30% sucrose solution. Coronal sections (50 *µ*m) were cut on a freezing microtome and stained (for cytochrome oxidase or for Nissl bodies) for microscopic verification of electrode positions.

Schematic outlines of brain structures and electrode tracks were drawn from photographs of histological slices and overlaid with brain atlas planes (GNU Image Manipulation Program, GIMP v 2.10, Inkscape v 1.0.1) to verify adequate placement of electrodes and estimate which probe channels recorded signal from cortex and which form the representation of large mystacial vibrissae located in dorsolateral part of VPM (Haidarliu et al., 2008) (Fig. 1 AB). Signals from these regions were used to estimate the similarity of raw and modeled EP (see below). Examples of histological data can be seen in the supplementary materials (Suppl. Fig. 2).

### 5.2 Stimulation and recordings

A group of large vibrissae from the left mystacial pad was glued to the piezoelectric stimulator at around 10–15 mm from the snout. Spike2 sequencer software (Cambridge Electronic Design) controlled square wave pulses of 1 or 2 ms duration and 20 V amplitude that produced a 0.1 mm horizontal (in the rostro-caudal axis) deflection of the whiskers. The stimuli (n=100) were delivered with 3–5 s pseudorandom intervals. Monopolar local field potentials (LFP) were recorded with multichannel silicon probes referenced to Ag/AgCl electrode placed under skin on the neck.

All cortical probes were tilted ∼30° from the vertical axis in order to enter the cortex roughly perpendicularly to its surface, within the barrel field (1.5–3.0 mm posterior and 5.0–6.2 right to bregma point). Thalamic electrodes were inserted vertically between 2.9–4.4 mm posterior to the bregma and between 2.3 and 3.7 mm lateral to middle-line. Several probe configurations were used to record evoked potentials (EP) from the thalamus and barrel cortex, whenever possible simultaneously, with multiple and dense arrays of electrodes. Probe models used for recordings included:

1. Single-shank Neuropixels (NP) v1.0 probes (Jun et al., 2017) with 384 recording channels (n=3 experiments). NP was configured into the “long column” pattern (https://billkarsh. github.io/SpikeGLX/#interesting-map-files) in which recording points span 7.68 mm of a tissue. NP recorded EP profile from all cortical layers, somatosensory thalamus and the intermediate tissue along the probe shank (Fig. 1).
2. Two eight-shank, 64 channels probes with 8x8 contact grid (0.2 mm inter-shank and inter-electrode distances, NeuroNexus A8x8), one of which was placed in the cortex, the other inserted into the thalamus (n=3 experiments). Each electrode grid covered 1.4 by 1.4 mm square within tissue. Whisker stimulation and EP recording were repeated with probes advanced to deeper locations (2 levels in the cortex and 3 levels around thalamus). Data were combined offline to form EP profiles covering the full depth of cortex (8x11 electrode grid) and a larger thalamic area (8x18 grid). For overlapping levels, data from only one penetration was included in the final dataset (example data can be seen in Fig. 3).
3. In the analysis we included also data from popular 16 channel probes: linear (A16) probes with 100 or 150 *µ*m spacing between contacts (NeuroNexus) and double shank probe (A2x8) with 1 mm inter-shank and 0.4 mm inter-electrode distance (ATLAS Neuroengineering). Two approaches were used: either one A16 probe was placed in the cortex and another A16 or A2x8 in the thalamus (n=2), or one linear probe was advanced obliquely from the cortex to the thalamus and recordings were repeated at consecutive depths (n=3). In the latter case, there was 200–600 *µ*m overlap (2–4 recording points) between consecutive probe positions. Similarly, as for the A8x8 probes, data from consecutive recording depths were combined to form longer EP profiles. For overlapping levels data from only one penetration was included in the final dataset.

Data from Neuropixels were filtered (0.5 Hz – 10 kHz), amplified (×50) and sampled (30 kHz) with SpikeGLX software (version 20190413, https://billkarsh.github.io/SpikeGLX/) to a binary data file. Stimuli triggers were recorded by an additional digital acquisition module (PXIe-6341, National Instruments) to a separate binary file.

For recordings with two NeuroNexus A8x8 probes we used 128-channel data acquisition (DAQ) system developed at the AGH University of Science and Technology in Cracow (Szypulska et al., 2016; Jurgielewicz et al., 2021). LFP signal was amplified (*×*500), filtered (0.6 Hz – 9 kHz) and sampled, along with stimuli triggers, at 40 kHz into HDF5 format files (The HDF Group, 2010).

The signal from pairs of 16-channel probes was amplified (×1000)) and filtered (0.3 Hz – 5 kHz) by two AM System 3600 amplifiers and sampled at 10 kHz with extended 32 channel CED 1401 power analog-digital interface. LFP was recorded along with the whisker stimulation markers by the Spike2 software (Cambridge Electronic Design) to a proprietary ^*^.smr data format files. Files were preprocessed using (Garcia et al., 2014).

### 5.3 Data preprocessing

Raw LFP data was loaded and analysed using *Numpy* and *Scipy* Python packages (Harris et al., 2020; Virtanen et al., 2020). Each of the three file formats used (SpikeGLX binary, HDF5 and smr) required dedicated algorithms for data import, preprocessing, trigger detection, and extraction of evoked potential sweeps. All scripts used to preprocess and analyse are available in a github repository (https://github.com/wsredniawa/LFP_reconstruction).

Data from all the experiments were additionally filtered and resampled to prepare uniform datasets with frequencies up to 2 kHz and with 5 kHz sampling rate. Signals around the stimulus (50 ms before and 100 ms after) were collected and averaged to represent evoked potentials (EPs) from all recording points. All channels were individually detrended by subtracting the mean amplitude from baseline level before stimulus. All presented results are based on average EPs.

For CSD analysis, it was necessary to define the spatial relationships between individual recording points. Therefore, histological outlines were aligned with atlas planes ((Paxinos and Watson, 2007), Suppl. Fig. 2C) to indicate stereotaxic coordinates of the probes deepest locations. These coordinates were then used to anchor electrode grids in a 3D model of the rat’s brain (Papp et al., 2014) according to patterns supplied by the producers. As mentioned above, the patterns were linear or rectangular with the exception of NP probes which recorded from a zig-zag grid with 20 *µ*m (vertical) and 16 *µ*m (horizontal) distance (see Fig. 1a in (Jun et al., 2017)). For kCSD model based reconstructions, CSD sources were homogeneously distributed around electrode locations inside cuboid shaped block. Additional 1 mm spacing was added in all directions to avoid edge effect of the reconstruction. For 1D visualization purposes (Fig. 2), CSDs were estimated and plotted at electrode locations. In NP, one channel is always dedicated to the reference and it cannot record the signal. In NeuroNexus probes, we typically had a few faulty electrodes. In case of the profiles combined from consecutive recording levels, we tried to have an overlap covering missing channels, but finally there were several missing points in all used probe setups. These data were excluded from the grids loaded for further CSD analyses.

### 5.4 Current source density reconstruction

We estimated the density of trans-membrane current sources by applying Kernel Current Source Density method (kCSD, Potworowski et al. (2012); Chintaluri et al. (2019, 2021)) to EPs (*V*) profiles. In this model-based method, we assume that the CSD profile *C* at *x* is spanned by a large set of *M* basis functions 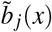 covering the region where we expect the activity to be estimated: 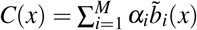. Each 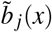 generates potential *b*_*j*_ (*x*) according to the assumed forward model taking into account specific geometry and conductivity of the system and forming a corresponding basis in the potential space: 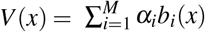. Conceptually, source estimation with kCSD is a two-step procedure. First, we do kernel interpolation of the measured potential using a symmetric kernel function 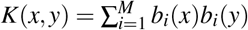. The regularized solution can be shown to be of the form (Potworowski et al., 2012) 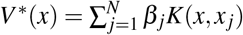, where *x* _*j*_ are electrode positions, *N* is their number, and *β* = (**K** + *λ* **I**)^*−*1^**V**, with **K**_*i, j*_ = *K*(*x*_*i*_, *x* _*j*_). The obtained solution must be translated to the CSD space which is achieved with a cross-kernel function 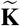, 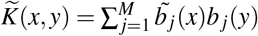, leading to the following CSD estimation:

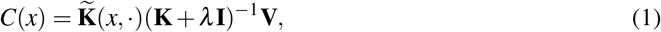

where
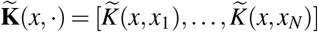. In the present paper we used three-dimensional gaussian functions as basis sources and assumed infinite tissue of constant conductivity. Regularization parameter *λ* was chosen with the L-curve regularization method (Chintaluri et al., 2019).

This approach, separating estimation space from the experimental setup, makes kernel CSD robust to missing nodes of EP profiles, while such data cannot be analyzed easily with classical CSD (Nicholson and Freeman, 1975).

Note that even if CSD basis sources 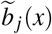 may in principle be well localized, each generates potential in the whole space. This allows modeling of various experimental situations in which electrodes are placed close or far from expected active current sources which we discussed in more detail previously (Chintaluri et al., 2021). In this paper we considered cortical only, thalamic only and cortico-thalamic sets of sources and electrode which were placed within the block of the rat’s brain model from (Papp et al., 2014) (see Fig. 3 and 5).

We used kCSD implementation available at https://github.com/Neuroinflab/kCSD-python, see (Chintaluri et al., 2019) for description. The analyses reported in this article were performed with a version from September 9, 2022.

### 5.5 Potential estimation from the subset of sources

To identify contributions to the LFP from a subregion *T*, say thalamus, let’s assume all basis sources 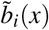 covering *T* are listed with *i* = 1, …, *L* while the rest are placed elsewhere (e.g. in the cortex). Then the total potential can be written as 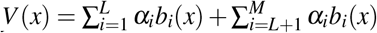. Clearly, the contribution from sources located in *T* is 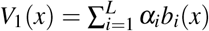. We can also split *K* into two parts defined by the basis functions in the two regions: 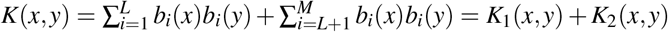. Since 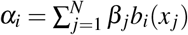, we can rephrase *V*_1_ as

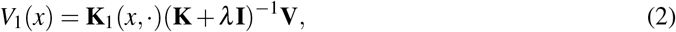

which, for *T* being thalamus, is the contribution of sources located within the thalamus to the total LFP.

### 5.6 Rolling correlation

To estimate similarity of EP waveforms we used rolling Pearson correlation score calculated in 3 ms overlapping windows with 1 ms step. Correlation coefficients were computed (1) between cortical and thalamic EPs (Fig. 1D), and (2) between estimated and measured EPs (Fig. 4). In the first case, we used a cortical channel from the middle cortical level, characterized by a large negative wave around 10 ms post stimulus and a thalamic recording from the center of the dorsal part of VPM nucleus (Fig. 1B–C). For a group analysis we averaged obtained coefficients across experiments (n=11) and represented them as a time series of mean correlation scores (e.g. Fig. 1D). Rolling correlation was used in a similar way to compare the time courses of measured EPs with their counterparts estimated from all the currents and from subsets of currents. The analysis was performed for EPs from the central locations in two structures separately.

### 5.7 Testing setups for effective/reliable thalamic CSD

Data from 8x8 NeuroNexus probes were used to model four experimental setups. We fed CSD algorithm with data from (1) only thalamic electrode grid or from (2) both thalamic and cortical grids; and we defined reconstruction space either (3) restricted to the main region of interest (thalamus or cortex) or (4) spanning the large volume including thalamic and cortical recording areas. We defined thalamic-only reconstruction space stretched on the 8x18 thalamic electrode grid with additional tissue margin of 0.5 mm in all 3 dimensions In the second variant, the space was extended dorsally and lateraly to a brain surface, and in an anterior direction to include all cortical recording point also with the margin 0.5 mm.

## 6 ACKNOWLEDGMENTS

The study received funding from the Polish National Science Centre’s grant 2013/08/W/NZ4/00691. The authors declare no conflict of interest.

Formatted in Overleaf with Basic Academic Journal Article Template (by John Hammersley CC BY 4.0)

