## supplementary materials for "Local contribution to the somatosensory evoked potentials in rat’s thalamus"

[1]Laboratory of Neurophysiology of Mind, Centre of Excellence for Neural Plasticity and Brain Disorders (BrainCity), Nencki Institute of Experimental Biology of Polish Academy of Sciences, 3 Pasteur Street, 02-093 Warsaw, Poland
[2]Neurobiology of Emotions Laboratory, Nencki Institute of Experimental Biology of Polish Academy of Sciences, 3 Pasteur Street, 02-093 Warsaw, Poland
[3]NeuroElectronics Research Flanders (imec, KU Leuven & VIB) Kapeldreef 75, 3001 Leuven, Belgium
[4]AGH University of Science and Technology in Kraków, Faculty of Physics and Applied Computer Science, al. Mickiewicza 30, 30-059 Krakow, Poland
[5]Laboratory of Neuroinformatics, Nencki Institute of Experimental Biology of Polish Academy of Sciences, 3 Pasteur Street, 02-093 Warsaw, Poland
[6]Jagiellonian University, Faculty of Management and Social Communication, Jagiellonian University, 30-348 Krakow, Poland

#### ABSTRACT

Local Field Potential (LFP), despite its name, often reflects remote activity. Depending on the orientation and synchrony of their sources, both oscillations and more complex waves may passively spread in brain tissue over long distances and be falsely interpreted as local activity at the recording site. Here, we study the contribution of local and distant currents to LFP recorded from rat thalamic nuclei and barrel cortex activated by whisker stimulation. We reconstructed the current sources using dense multichannel recordings and a model-based kernel Current Source Density (kCSD) method. We show that the evoked potential wave seen in the thalamic nuclei around 7–15 ms post-stimulus has a substantial negative component reaching from cortex. This component can be analytically removed and truly local thalamic LFP, with purely thalamic contributions, can be recovered reliably using kCSD. In particular, concurrent recordings from the cortex are not essential for reliable thalamic CSD estimation. Proposed framework can be used to analyse LFP from other brain areas and has consequences for general LFP interpretation and analysis.

Keywords: subcortical signal, field potentials, volume conduction, CSD, vibrissa-barrel system

### 1 SUPPLEMENTARY RESULTS

#### 1.1 Pharmacological silencing of the barrel cortex

To additionally validate our conclusions regarding the input of cortical field potentials to thalamic recording, in one experiment (rat NP-2) we reduced cortical activity with topical application of a sodium channel blocker lidocaine (Lignocainum Hydrochloricum 2% Polfa Warszawa). Whisker evoked responses were recorded in a control, physiological condition (this data is included in a main analyzed dataset) and 60 min after lidocaine perfusion. As expected, lidocaine substantially reduced cortical activity and practically erased large waves of whisker evoked cortical responses (compare Suppl. Fig. 1A with B). There was no such evident change in the thalamus, where early multiunit activity was equally intensive before and after lidocaine. We applied our analysis pipeline (recorded LFP -¿ kCSD -¿ LFP reconstruction from a thalamic subset of sources) to thalamic EPs recorded control and post-lidocaine condition. In a control condition, there was a clear difference between recorded and reconstructed EPs (Suppl. Fig. 1D), reaching a statistical significance around 10 ms post stimulus (Suppl. Fig. 1E). This difference, which we believe results from cortical impact, indeed disappeared after lidocaine application. Recorded and reconstructed EPs in lidocaine conditions are practically identical (Suppl. Fig. 1D, E; red lines). Additionally, we extracted multiunit action potentials (combinato package, Niediek et al. (2016)) from the same thalamic channel as shown in panel D.

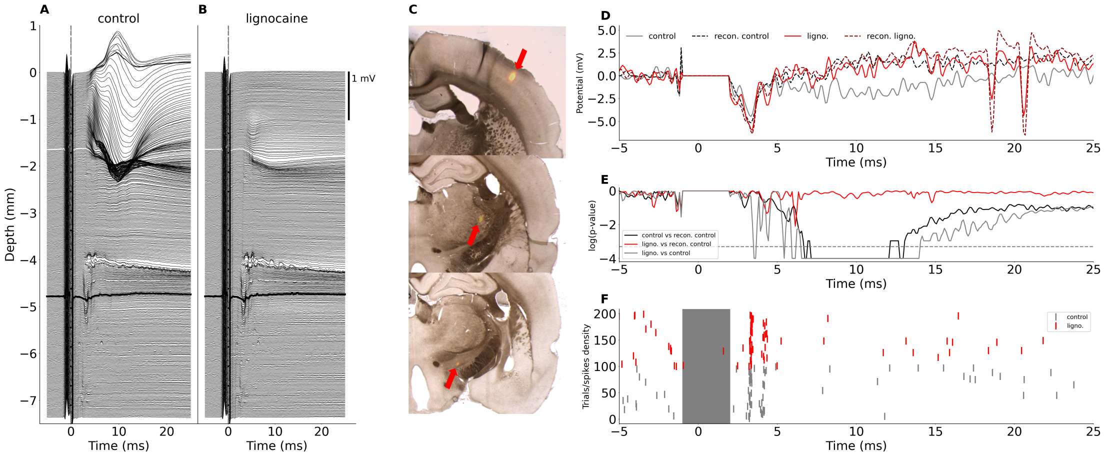


**Supplementary Figure 1.** Experiment NP-2. A and B: EP traces obtained from recording points along Neuropixels probe in control condition (A) and 60 minutes after lidocaine perfusion (B). Thick lines in A and B indicate EP traces further analyzed in panels D and E. C: Trace of Neuropixels probe in the histological brain slices from lidocaine experiment. D: Overlay of measured and estimated thalamic EPs (single trial) from control and lidocaine condition. Continuous lines represent real EPs, dashed lines — reconstructions. Note that without the influence of cortical field eliminated by lidocaine, recorded thalamic potential became more positive, just like suggested by the CSD-based estimation. E: p-values for the statistical comparison (permutation test) of recorded and reconstructed EP traces: black line — control condition; red line — post-lidocaine condition). Gray line plots p values for the comparison of control versus post-lidocaine EP reconstructions. Note, that p-values are plotted in the logarithmic scale, horizontal dashed line marks the level of P=0.05. F: spiking activity (raster plot) from the indicated channel in the thalamus in control (gray) and after lidocaine (red). Grey shading shows region of artifact removal.

#### 1.2 Histology

Schematic outlines of brain structures and electrode tracks were drawn from photographs of histological slices (viewed in white and green for a DiI fluorescence light) and overlaid with brain atlas planes (GNU Image Manipulation Program, GIMP v 2.10, Inkscape v 1.0.1) to verify adequate placement of electrodes and estimate which probe channels recorded signal from cortical (barrel field) and which from the thalamic representation of large mystacial vibrissae (dorsolateral part of VPM).

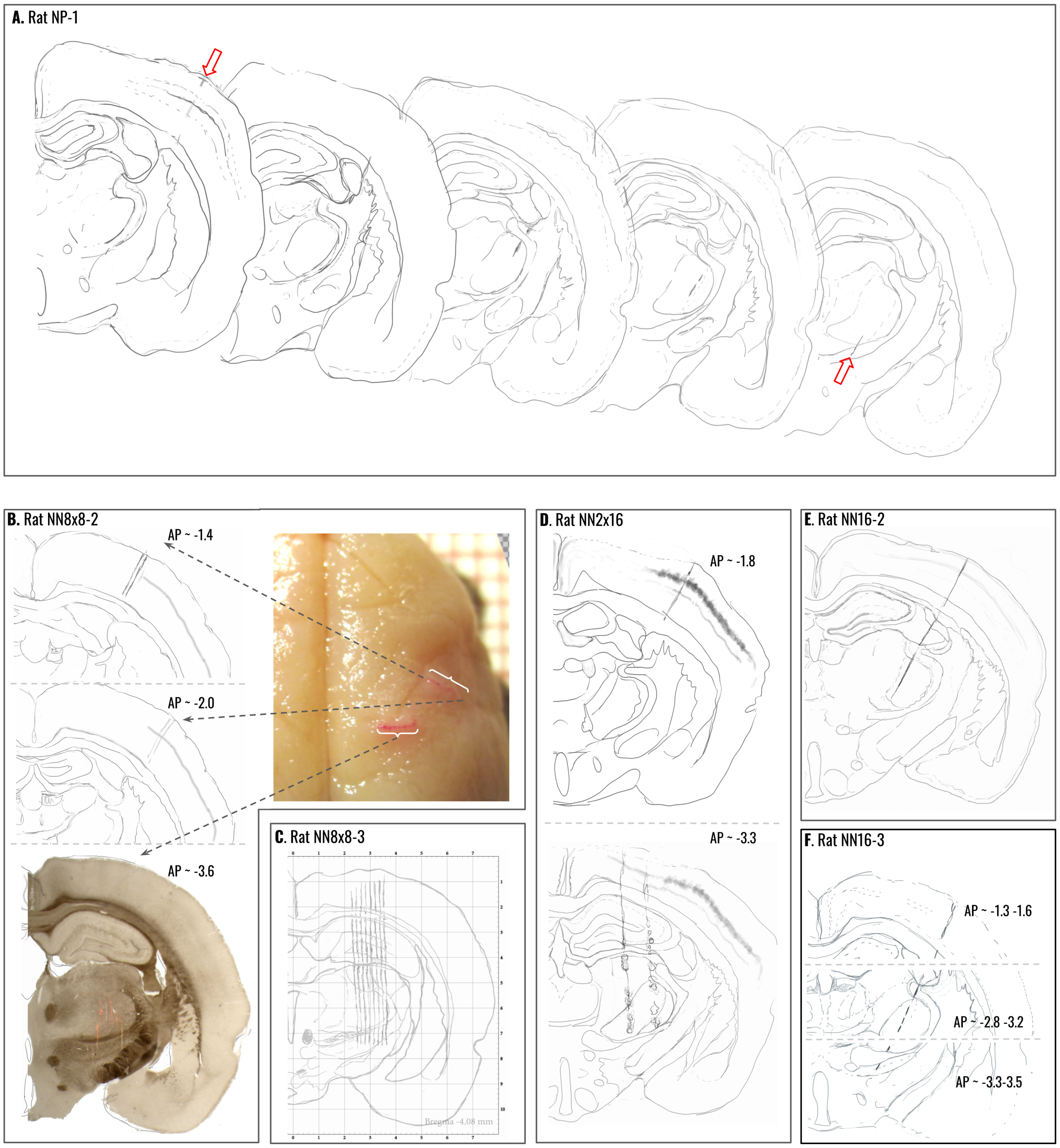


**Supplementary Figure 2.** Examples of histological verification of electrodes' locations. See text below for description.

Consecutive panels in Suppl. Fig. 2 present examples from different electrode setups. A: Typical histology from an experiment with Neuropixels probe penetrating right hemisphere from brain surface in the barrel cortex obliquely down, medial and posterior towards deep thalamic location. Outlines of consecutive coronal slices following electrode trace. See also Suppl. Fig. 1 for an example from another Neuropixels experiment. B: A simultaneous recording with two 2D (8x8) NeuroNexus silicone probes placed in the cortex and the thalamus. Entrances of both probes are evident in the top view of the fixed brain. Left upper panels show outlines of two slices with the most anterior and most posterior probe shank traces in the barrel cortex. The lower panel presents a photograph of a wet, unstained slice with DiI traces of electrodes penetrating the thalamus. C: Another example with NeuroNexus A8x8 probe in thalamus. Outline of a brain slice is overlaid on 1x1 mm atlas grid to facilitate estimation of electrode positions in stereotaxic coordinates. D: Simultaneous recording with a linear (1x16) NeuroNexus silicone probe in the barrel cortex and 2x8 (Atlas) probe in the thalamus. E-F: Electrode traces left in tissue by a single NeuroNexus 1x16 probe advanced obliquely from barrel cortex to deeper and more medial thalamic sites. In E, the trace was almost completely visualised within a single slice, while in F, because of larger electrode anterior-posterior tilt, the trace was visible through multiple consecutive slices. Histological outlines were used to choose the recording points located centrally within cortical and thalamic whisker representation. Data from these channels were used for the group analyses and visualization of results.

##### 1.3 Statistical assessment of veracity of CSD estimation in four considered scenarios

Our experiment verified that LFP signal recorded in the thalamus after whisker stimulation is built up by contributions from weak local currents and by passively conducted contributions from strong cortical activation. To verify the necessity for inclusion of remote electrodes on reliable estimation of thalamic currents we compared four variants of electrodes' locations and reconstruction spaces (see Fig. 5 in the main text). Here, the most extensive scenario (with grids of electrodes and CSD bases both in thalamus and barrel cortex, Fig. 5C) was considered a reference. Veracity of reconstruction in the other cases was assessed by analyzing differences of reconstructed CSD profiles from the reference one with the following metrics (Chintaluri et al. (2019)): $\mathrm{err}(x) = |\tilde{C(x)}/\|\tilde{C(x)}\| - C(x)/\|C(x)\||$, where $C(x)$ is the reference and $\tilde{C(x)}$ is an alternative reconstruction. We used $L_2$ normalization.

To evaluate statistical significance of obtained differences we performed permutation tests for each pixel of the CSD map. Each single trial (N = 100) was analysed using each reconstruction subversion giving a total of 400 CSD maps (for each time point). To compute the p-value maps, real metric's ($err(x)$) score was used as a threshold for metrics' distribution obtained with permutations (N=1000). The analysis was run separately for three analysed time points (5, 10, 20 ms poststimulus).

Supplementary Fig. 3 below presents the maps of the probability that for a given setup, CSD estimation is not different from the reference one (i.e. the reconstruction subversion is equally valid as the reference one). The p-value map for the first solution (Supp. Fig 3A) shows no similarity to the reference map in the whole space as this setup for CSD reconstruction as incorrect. For two other alternative solutions (Supp. Fig 3C and 3D) p-value maps show high similarity in the subcortical space. Differences in cortical area are obvious as we are not using information from cortical channels for alternative approaches. However, there are some spots around/within thlamic nuclei that differ between reconstructions mostly at the timepoint when both, cortex and thalamus, are active (around 10 ms after onset).

#### 2 SUPPLEMENTARY METHODS

##### 2.1 Data preprocessing

Raw LFP data were loaded and analysed using *Numpy* and *Scipy* Python packages. Each of the three file formats used to store acquired data — SpikeGLX binary, HDF5, and .smr — required dedicated algorithms for data import and preprocessing, trigger detection, and extraction of evoked potential sweeps. Here we include additional information on these preprocessing steps.

###### *2.1.1 NeuroPixels (NP) data*

For NP data stimuli triggers were recorded in separate binary files. They were off-line synchronized with the Neuropixels' clock signal using CatGT and TPrime software (https://billkarsh.github.io/SpikeGLX/catgt).

###### *2.1.2 HDF5 files*

HDF5 files from AGH DAQ included LFP and trigger channels that were all sampled simultaneously, with no inter-channel delays, which allowed easy EP detection and extraction from continuous data. Additional headstage-to-probe channels mapping and electrode layout files allowed to reconstruct spatial arrangement of data from all recorded channels.

###### *2.1.3 Spike2 data format*

Files in Spike2 .smr format also included digital channels with stimulus markers that were sampled simultaneously with LFP, which allowed easy EP detection and extraction from continuous data. Similarly as for A8x8 probes, data from consecutive recording depths were combined to form longer EP profiles.

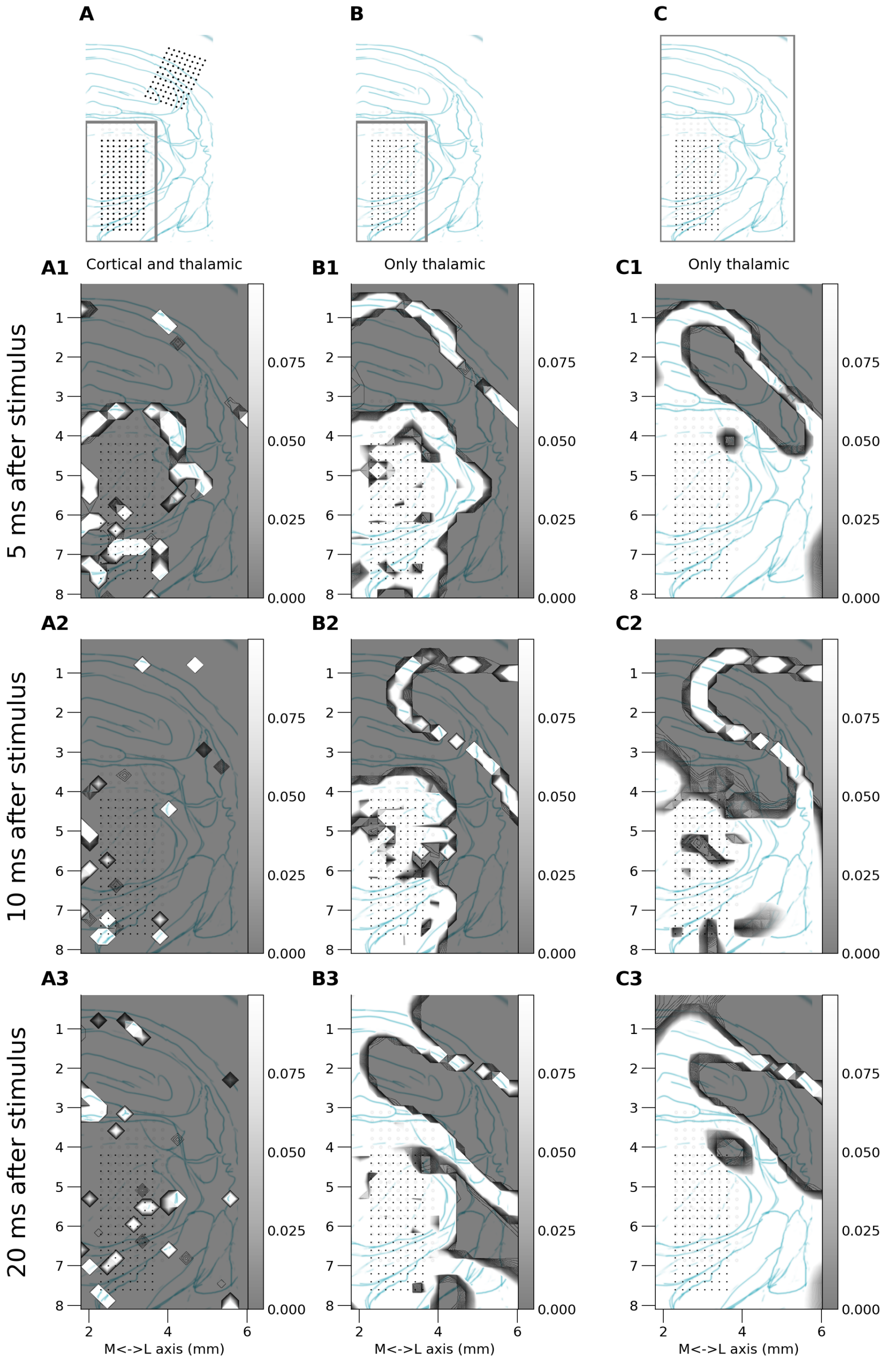


**Supplementary Figure 3.** Statistical comparison of the alternative approaches to CSD reconstruction (compare Fig. 5 of the article). Shading represents statistical significance of metrics between subversion of reconstruction and the full one (Fig. 5C). Upper row schematically indicates the space occupied by considered electrodes and sources. A: Recordings from the thalamus and cortex considered; sources assumed only in the thalamus. B: Only thalamic recordings considered; sources assumed only in the thalamus. C: Recordings from the thalamus; reconstruction in a large block of tissue covering thalamus and cortex. Rows 1–3 show CSD reconstructions snapshots at 5, 10 and 20 ms after stimulus that were analyzed assuming setups indicated in the top row. P-value spatial maps are overlaid on a drawing representing histological verification of recording points (dots) location within thalamic structures. p-value score for every pixel is coded with gray color. The darker the area the less we can trust the alternative reconstruction.

###### 2.1.4 Locations of recording points used in consecutive analyses

Suppl. Table 1 lists electrode types and numbers of data channels used for group analyses and illustrations. NN16 — one shaft, 16 channels Neuronexus silicone probe; Atlas2x8 — two shaft, 16 channels Atlas silicone probe; NN8x8 — 8 shaft, 64 channels Neuronexus silicone probe; NP — Neuropixels v.1 probe. In case of one-probe setups, a single probe was inserted obliquely from cortex to thalamus, otherwise, one probe was inserted to cortex the other vertically to thalamus. NP probes recorded both structures in one oblique penetration. All other shorter probes were advanced deeper and deeper into the tissue and recordings were made at consecutive levels. Data from all electrodes and all levels were combined into a single dataset covering all the recording space. Indicated numbers refers to the channels in such full datasets from each experiment.

**Supplementary Table 1.** Electrode(s) type and numbers of data channels used for group analyses and illustrations

| | experiment ID | probe(s) type | channel No - cortex | channel No - thalamus |
|---|---|---|---|---|
| 0 | NN16-1 | 1 x NN16 | 6 | 28 |
| 1 | NN16-2 | 1 x NN16 | 7 | 27 |
| 2 | NN16-3 | 1 x NN16 | 8 | 26 |
| 3 | NN16-4 | 1 x NN16 | 8 | 34 |
| 4 | NN2x16 | 1 x NN16 + 1 x Atlas2x8 | 6 | 15 |
| 5 | NN8x8-1 | 2 x NN8x8 | 60 | 114 |
| 6 | NN8x8-2 | 2 x NN8x8 | 78 | 158 |
| 7 | NN8x8-3 | 2 x NN8x8 | 78 | 152 |
| 8 | NP-1 | 1 x NP | 320 | 124 |
| 9 | NP-2 | 1 x NP | 330 | 134 |
| 10 | NP-3 | 1 x NP | 320 | 130 |

### 3 SCRIPTS USED TO PREPROCESS AND ANALYZE DATA

All scripts to preprocess raw data and then compute CSD and reconstruct LFP are available in a github repository (https://github.com/wsredniawa/LFP_recon). We can provide raw data on demand. Please, contact us at the corresponding authors email addresses.
